# Plant inositol transport influences bacterial colonization phenotypes

**DOI:** 10.1101/2023.05.03.538913

**Authors:** Bridget S. O’Banion, Piet Jones, Alexander A. Demetros, Brittni R. Kelley, Andrew S. Wagner, Jin-Gui Chen, Wellington Muchero, Todd B. Reynolds, Daniel Jacobson, Sarah L. Lebeis

**Affiliations:** Department of Microbiology, University of Tennessee, Knoxville, TN 37996, USA; The Bredesen Center for Interdisciplinary Research and Graduate Education, University of Tennessee, Knoxville, TN 37996, USA; Department of Microbiology and Molecular Genetics, Michigan State University, East Lansing, MI 48824, USA; Plant Resilience Institute, Michigan State University, East Lansing, MI 48824, USA; Biosciences, Oak Ridge National Laboratory, TN 37830, USA; Department of Plant, Soil and Microbial Sciences, Michigan State University, East Lansing, MI 48824, USA

**Keywords:** Host-microbe interactions, inositol, root colonization, GWAS, Arabidopsis

## Abstract

Plant microbiomes are assembled and modified through a complex milieu of biotic and abiotic factors. Despite dynamic and fluctuating contributing variables, specific host metabolites are consistently identified as important mediators of microbial interactions. We combine information from a large-scale metatranscriptomic dataset from natural poplar trees and experimental genetic manipulation assays in model Arabidopsis seedlings to converge on a conserved role for transport of the plant metabolite *myo*-inositol in mediating host-microbe interactions. While microbial catabolism of this compound is often linked to increased host colonization, we identify motility phenotypes that occur independently of catabolism, suggesting that inositol may additionally serve as a eukaryotic-derived signaling molecule to modulate microbial activities. Our data suggests host control of this compound and resulting microbial behavior are important mechanisms at play surrounding the host metabolite inositol.

## Introduction

Plants and microorganisms use a variety of mechanisms to establish and maintain complex interkingdom interactions at the root-soil interface ^1–3^. As plants dynamically release nutrient-rich root exudates into the surrounding soil, the structure of their rhizosphere microbiome responds via shifts in total abundance, composition, and activity ^4,5^. A portion of this community subsequently colonizes the external rhizoplane and internal endophytic root tissue, which requires increasingly intimate interactions with the physical root structure, host immune response, and additional microbiome members ^5–7^. Many research efforts have focused on disentangling the processes mediating initial bacterial attraction and attachment to the root surface, including chemotaxis and biofilm formation ^3,8–10^. However, less is known about mechanisms broadly employed by both plants and microbes to maintain these interactions at the root interface over time.

Diverse experimental approaches suggest plant-associated microorganisms enhance various facets of plant health by improving nutrient acquisition, biomass, and resistance to biotic and abiotic stressors ^11^. Large sequencing and analytical datasets (*i.e.*, ‘-omics studies) provide researchers with abundant data to parse for influencing factors, such as genes or metabolites. While these types of studies are often conducted under conditions that more accurately represent the complexity of natural systems, it is difficult to explicitly demonstrate causative mechanisms ^12^. Alternatively, laboratory experiments using axenic systems allow a thorough investigation of physiological intricacies but can be difficult to translate to *in situ* conditions ^13^. Potential mechanisms identified across these contrasting approaches represent attractive targets for further research, as their biological signal is observed under various conditions ^14^.

Plant metabolites are well-documented mediators of cross-kingdom interactions across a multitude of experimental designs ^4,14^. For example, root exudates change in response to pathogen infection, developmental stage, abiotic stress, and more ^4,15–17^. These altered exudate patterns also correlate with changing microbial communities, revealing host chemical manipulation of the rhizosphere microbiome ^15,18^. Importantly, dynamic root exudate composition suggests these compounds are also differentially produced and trafficked through plant tissue and likely influence rhizoplane and endophytic microorganisms as well. While plants can synthesize a variety of antimicrobial compounds ^19^, many common root exudate components, such as sugars and organic acids, can recruit and repel microorganisms via chemotaxis and serve as nutrient sources ^4,10,20^. However, the role these compounds may play beyond initial recruitment and providing a labile rhizosphere nutrient source has not been investigated as thoroughly, especially relating to non-pathogenic and non-beneficial microbial strains.

Here, we describe the contribution of a host-derived metabolite to bacterial root colonization outcomes. We use information from a large metatranscriptomic dataset to infer a role for the plant metabolite *myo*-inositol in the relationship between plant hosts and their microbiome. To explore this observation further, we employed a suite of host and bacterial mutant lines, as well as diverse bacterial isolates, to investigate host and microbial genetic factors that contribute to observed colonization patterns. By linking these varied experimental approaches, we shed light on a conserved *myo*-inositol dependent mechanism underpinning plant-microbe interactions across diverse plant and microbial phylogenies, environmental conditions, and community complexity.

## Results

### GWAS sub-networks identify putative poplar genes influencing diverse microbial interactions

To identify host mechanisms influencing microbial dynamics in a complex environment, we used a metatranscriptome dataset^21^ to integrate microbial taxa and host genome variant information from field-grown *Populus trichocarpa* (poplar) trees. A genome-wide association study (GWAS) was performed to identify host single nucleotide polymorphisms (SNPs) that significantly correlate (p-value < 1e-5) with the pseudo-abundance of microbial transcripts. We created GWAS sub-networks focused on transcripts mapping to two microbial genera, *Pantoea* and *Streptomyces*, which are both cosmopolitan plant microbiome members from distinct phyla (Proteobacteria and Actinobacteria, respectively). We hypothesized that these analyses would identify SNPs in distinct host genes correlating with the colonization dynamics of diverse microbial constituents.

The sub-networks identified many SNPs across diverse poplar genes that correlated with transcript levels of our genera of interest (Figure 1A, Dataset S1). Although sub-networks were generated from both xylem and leaf tissue datasets, the leaf tissue provided no significant results for the *Pantoea* sub-network, and only 7 SNPs across 4 poplar genes for the *Streptomyces* sub-network. Therefore, we pursued the xylem-based networks for further insights. In the *Pantoea*-specific xylem dataset, 13 SNPs were identified across 8 poplar genes. Alternatively, the *Streptomyces*-specific dataset identified 76 SNPs across 49 genes (Figure 1A). All poplar genes were annotated with mercator to assign a MapMan ontology and identify the best hits to the *Arabidopsis thaliana* (Arabidopsis) reference database ^22^. The identified genes in the *Pantoea* sub-network are involved in various biological processes, including transcriptional regulation, carbohydrate metabolism, and transport. The *Streptomyces* sub-network revealed host genes involved in abiotic and biotic stress signaling, protein synthesis and degradation, transport, and more (Figure 1A). Together, these GWAS networks suggest a role for diverse poplar genes in mediating the activity of evolutionarily distinct microbiome members.

**Figure 1.**
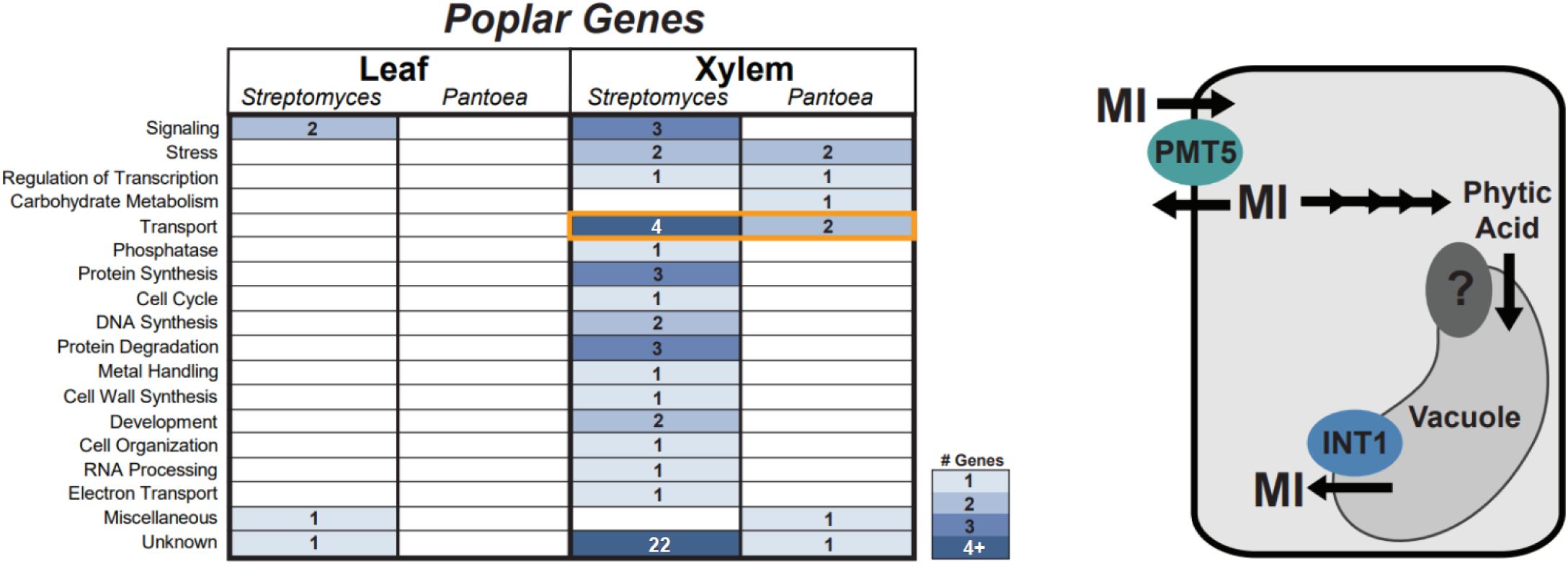
GWAS sub-networks identify putative poplar genes influencing diverse microbial interactions. Meta-transcriptomic sequencing data from *Populus trichocarpa* leaf and xylem tissue was used to generate GWAS sub-networks that identified plant SNPs correlated to transcript pseudo-abundances for the microbial genera *Streptomyces* and *Pantoea*. A) The resulting sub-networks identified diverse genes that may influence colonization dynamics in the leaf and xylem tissue. Categories are labeled based on the associated MapMan description (Dataset S1). The number of identified genes within each category are labeled and shaded in blue. White boxes indicate no gene hits. Both xylem sub-networks contain genes in the Transport category (orange box) with high sequence identity to previously characterized Arabidopsis inositol transporters. B) A simplified diagram showing the predicted cellular localization of the Arabidopsis homologs of the two transporters identified by the poplar GWAS. *Myo*-inositol (MI) is transported across the plasma membrane by PMT5 (teal ellipse) and across the tonoplast by INT1 (blue ellipse). Vacuolar inositol is a product of phytic acid dephosphorylation. Soluble inositol can be phosphorylated into phytic acid before transport into the vacuole (light gray shape) via an uncharacterized transporter (dark gray ellipse).

Intriguingly, we found finer-scale functional redundancy among the poplar genes identified by the two sub-networks. A single host gene with one SNP in the *Streptomyces* network, and two poplar genes with three total SNPs in the *Pantoea* network, mapped to Arabidopsis genes characterized as *myo*-inositol transporters (AT2G43330, *INT1*; AT3G18830, *PMT5*) (Figures 1A and 1B). Inositol is a cyclic polyol known to serve as a critical eukaryotic structural and signaling molecule ^23^. The most abundant inositol isomer, *myo*-inositol (herein referred to as inositol), is found in root exudates ^4^ and has been implicated in various beneficial and pathogenic plant-microbe interactions ^24,25^. We therefore chose to directly investigate the role of the identified host genes by pivoting to laboratory experiments using Arabidopsis.

### Arabidopsis inositol transporters mediate microbial root colonization levels

To directly interrogate the role of host inositol transport on microbial colonization dynamics, we obtained Arabidopsis genotypes with T-DNA insertions in the genes of interest. The gene identified in the *Pantoea* sub-network, *INT1*, encodes a tonoplast-localized transporter that specifically exports vacuolar inositol to the cytoplasm (Figure 1B) ^26^. Alternatively, the *Streptomyces* sub-network identified a poplar homolog of *PMT5*, which encodes a transporter shown to move various polyols and monosaccharides, including inositol, across the plasma membrane of Arabidopsis (Figure 1B) ^27^. We then performed colonization assays pairing a representative *Pantoea* and *Streptomyces* isolate (Table S1) with seedlings of the corresponding Arabidopsis T-DNA insertion lines (*int1* for *Pantoea*; *pmt5* for *Streptomyces*). After 7 days, roots were weighed, washed, and homogenized to plate for viable bacteria. Because the host transporters were not identified in the poplar leaf-centric GWAS sub-networks and root expression of each transporter has been previously confirmed, we focused specifically on Arabidopsis root tissue ^26,27^. Intriguingly, *Streptomyces sp*. CL18 and *Pantoea sp*. R4 colonized the roots of *pmt5* and *int1*, respectively, at significantly lower levels than Arabidopsis Col-0 accession (Figures 2A and 2B).

**Figure 2.**
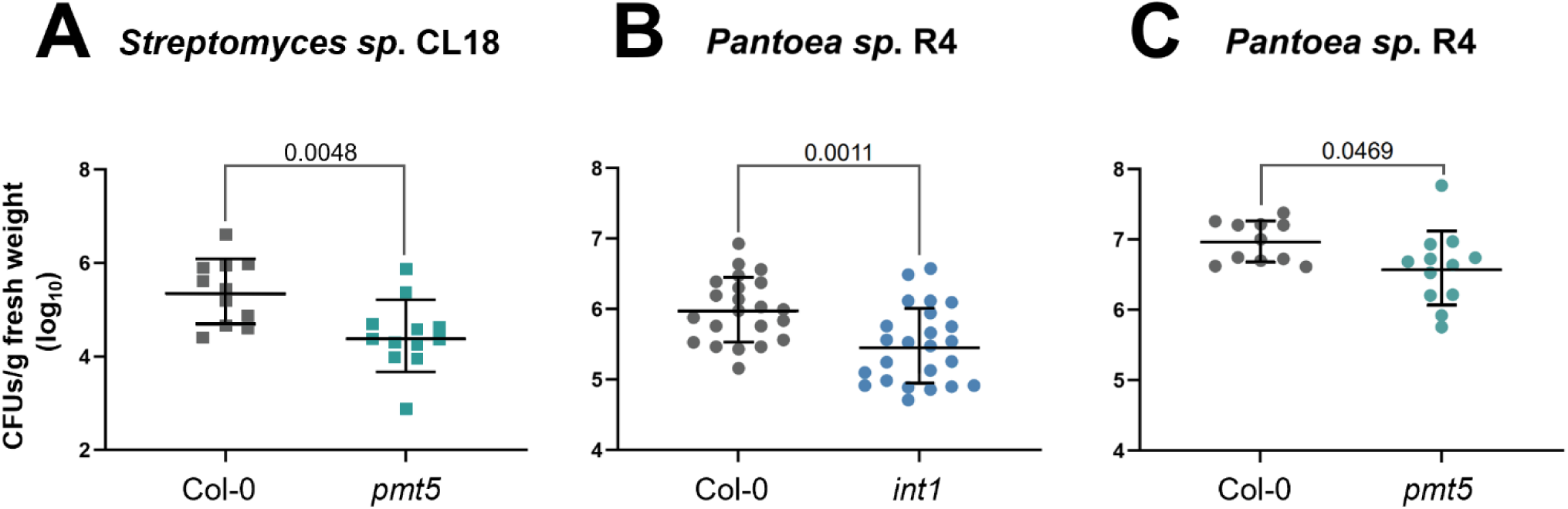
Arabidopsis inositol transporters mediate microbial root colonization levels. To directly investigate the role of inositol transporters in mediating microbial interactions, we performed 7-day colonization assays using representative bacterial isolates and Arabidopsis T-DNA insertion lines (*pmt5* and *int1*). A) *Streptomyces sp*. CL18 (square symbols) colonization of Col-0 (gray) and *pmt5* (teal) roots. *Pantoea sp*. R4 (circular symbols) colonization of Col-0 (gray) and *int1* (B; blue) or *pmt5* (C; teal). In all panels, each symbol represents the pooled roots of 3 seedlings. Data points from all panels fit a lognormal distribution and were therefore log-transformed prior to performing an unpaired t-test with Welch’s correction. Significant p-values (< 0.05) are displayed on each graph. Bars indicate the geometric mean +/- geometric standard deviation (SD).

Because PMT5 transports a variety of polyols and monosaccharides across the plasma membrane ^27^, the observed *Streptomyces* colonization phenotype could be due to altered dynamics of other sugar molecules within the host. We therefore repeated colonization assays with the addition of exogenous inositol to the media. Importantly, the root colonization deficit was rescued upon inositol addition to both the *Streptomyces*-*pmt5* and *Pantoea*-*int1* colonization assays (Figures S1A and S1B), suggesting this metabolite is specifically and sufficiently able to mediate the observed phenotype. It is also important to note that root colonization levels do not increase when inositol is added to Col-0 assays, suggesting this compound is not just providing a general growth advantage to microorganisms intimately associated with host tissue (Figures S1C and S1D).

We next investigated the role of additional enzymes in the host inositol pathway on microbial colonization. Because 25% of *Pantoea* sub-network host genes mapped to inositol transporters (Figure 1A), we focused specifically on *Pantoea* colonization dynamics moving forward. We characterized the colonization levels of *Pantoea sp*. R4 on the host transporter mutant identified by the *Streptomyces* sub-network (*pmt5*), as well as T-DNA insertion lines in a *myo*-inositol phosphate synthase (*mips2*) and a downstream inositol phosphate kinase (*ipk1*). These enzymes perform critical steps in plant inositol biosynthesis and transformation (Figure S2A) ^23^. *Pantoea* root colonization levels were significantly reduced on *pmt5* seedlings compared to Col-0 (Figure 2C). This is notable, considering this transporter was identified by the *Streptomyces* sub-network and controls metabolite flux at a different cellular location compared to INT1 (Figure 1B). Exogenous inositol similarly rescued *Pantoea sp*. R4’s colonization deficit on this promiscuous sugar transporter mutant (Fig S1E). Alternatively, colonization levels on *mips2* and *ipk1* seedlings were not significantly different from Col-0 levels, regardless of inositol supplementation (Figures S2B and S2C). Together, these experimental assays validate the importance of putative host genes identified by computational approaches and define a conserved role for host inositol transport in mediating colonization across diverse hosts (poplar and Arabidopsis) and bacterial phyla (Actinobacteria and Proteobacteria).

### Proteobacterial inositol catabolism correlates with higher root colonization

While inositol is integral to various eukaryotic cellular processes, bacteria generally use it primarily as a carbon source, with few known exceptions ^28–30^. Both *Pantoea sp*. R4 and *Streptomyces sp*. CL18 encode homologs of a previously characterized pathway for inositol catabolism ^28^ and grow in minimal media containing inositol as a sole carbon source (Figure S3A-C). Therefore, the lowered colonization levels in the *pmt5* and *int1* lines may be due to the bacterium losing access to a host-derived carbon source. To broadly investigate the relationship between bacterial inositol catabolism and host colonization, we analyzed the root colonization levels of a diverse set of proteobacterial isolates (Figure 3A) with varying abilities to use inositol as a sole carbon source (Tables S1 and S2). We hypothesized that microorganisms capable of growing on inositol would colonize Arabidopsis at higher levels than those unable to utilize this compound.

**Figure 3.**
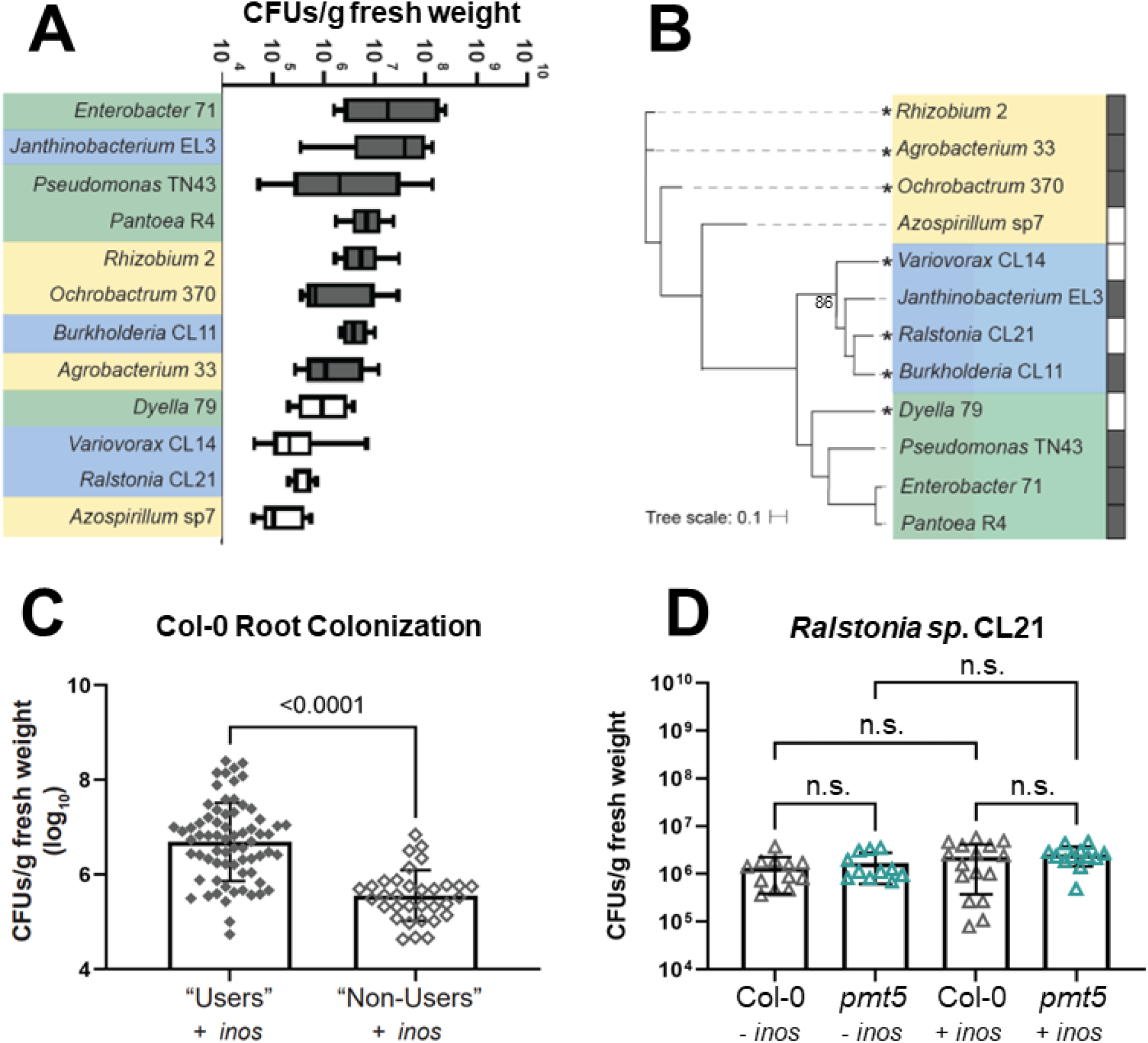
Proteobacterial inositol catabolism correlates with higher root colonization. We used a collection of diverse bacterial isolates to further connect bacterial inositol catabolism to host colonization levels. The isolates were associated with Col-0 seedlings in magenta jars (MS media + inositol). Each jar contained 5 seedlings. After 2 weeks, the 5 seedlings from each jar were pooled and bacterial CFUs in the root were enumerated (n=8-11 jars). A) The root colonization levels of 12 proteobacterial isolates are ordered from highest to lowest average CFUs. Isolates that can catabolize inositol have gray box-and-whisker plots (“users”), while those that cannot use inositol as a sole carbon source have white plots (“non-users”). Whiskers represent min to max values. Colored shading mimics taxonomic coloring in panel B. B) A phylogenetic tree built using a set of 27 concatenated genes showcases the relatedness of the twelve isolates used in this assay. Alphaproteobacteria (yellow shading), betaproteobacteria (blue shading), and gammaproteobacteria (green shading) cluster distinctly. Tree branches with an asterisk indicate strains that were originally isolated from Arabidopsis tissue. Dark gray rectangles next to isolate names indicate those that can use inositol as a sole carbon source (white boxes are unable to use this compound, as in the box-and-whisker plots in panel A). Bootstrap values are 100 unless otherwise indicated. Panel C shows the individual data points from panel A, aggregated into binary groups of inositol “users” (filled diamond symbols) and “non-users” (open diamond symbols). Each symbol represents pooled roots of 5 seedlings. Data points in panel C fit a lognormal distribution and were therefore log-transformed prior to performing an unpaired t-test with Welch’s correction. D) A selected proteobacterial isolate that cannot use inositol as a sole carbon source, *Ralstonia* CL21 (open triangle symbols), was inoculated onto Col-0 (gray) and *pmt5* (teal) plants. Each symbol represents the pooled roots of 3 seedlings. Data points in panel D were analyzed with a Kruskal—Wallis and Dunn’s multiple comparison test. Significant p-values (< 0.05) are shown, and bars represent the mean +/- SD. Inositol treatment (+/- inos) is indicated in panels C and D.

We measured root colonization levels on Arabidopsis Col-0 for 12 isolates representing 10 families commonly found in plant microbiomes (*e.g.,* Burkholderiaceae, Rhizobiaceae, and Pseudomonadaceae; Table S1) ^7,31^. These strains were originally isolated from various plants or soil and have fully sequenced genomes. Of the 12 isolates, 8 encode a putative genetic pathway for inositol catabolism (Figure 3B). We performed growth curves to verify these pathways are functional and confer the ability to use inositol as a sole carbon source (Table S2). Intriguingly, when the isolates are ordered from highest to lowest average CFUs/gram biomass, the 8 strains capable of inositol catabolism are first, followed by those that cannot use this carbon source (Figure 3A). In fact, when root colonization data-points are aggregated into binary groups (inositol “users” vs. “non-users”), the “users” colonize roots at an average of 31.1x higher cell numbers than “non-users” (Figure 3C). Additionally, neither colonization level nor inositol catabolism seem to align with microbial phylogeny or the original plant host (Figure 3A and 3B). We also measured the cell numbers for these isolates in the rhizosphere and bulk media of our experimental setup and observed that inositol catabolism patterns did not match the highest and lowest colonizers in these fractions (Figures S4A and S4B). Further, while “users” colonized the rhizosphere and bulk media at significantly higher levels than “non-users”, the magnitude is much more modest (1.6 and 3.9x higher, respectively). These proteobacterial assays were conducted in media containing inositol, and it is therefore not unexpected for this carbon source to provide a growth advantage to “users” living outside of the plant. This makes the dramatic increase in root colonization levels more intriguing, and lends further support for this compound mediating intimate associations with the root tissue specifically.

While the genetic and phenotypic potential for inositol catabolism seems to correlate with higher root colonization, there could be many other factors contributing to this phenotype across our proteobacterial collection. We thus compared the genomes of the 8 highest colonizers (“users”) to those of the lowest 4 (“non-users”) to identify other potential genetic determinants of elevated root colonization. Impressively, via two separate computational approaches, genes in the inositol catabolism pathway are the majority of the distinguishing genetic factors between these groups (Table S3). In addition to inositol catabolism genes, these approaches each identified an additional transporter as genetically distinct between these groups. While this comparative genomic approach does not consider regulation of these genes *in vivo*, it is notable that a catabolic pathway for a plant-derived metabolite is genetically indicative of elevated root colonization levels among our bacterial strains.

We were next curious whether the colonization deficits observed on the Arabidopsis transporter mutants were specific to microorganisms capable of catabolizing inositol or were more general effects due to deleterious impacts on host physiology. Therefore, we selected one of the proteobacterial isolates described above that is unable to use inositol as a sole carbon source, *Ralstonia* CL21, for additional colonization assays with *pmt5* seedlings. After 1 week, there was no difference in the root colonization levels between Col-0 and *pmt5* genotypes, regardless of exogenous inositol addition (Figure 3D). Root colonization on *pmt5* is therefore uniquely impacted only for two tested isolates with the inositol catabolic pathway (*Streptomyces sp.* CL18 and *Pantoea sp*. R4; Figure 2A and C). Altogether, these results provide correlative evidence of a relationship between root colonization phenotypes and microbial inositol catabolism.

### Pantoea sp. R4 inositol catabolism does not facilitate increased root colonization

While interrupting host transport of inositol impacts the colonization of diverse bacterial isolates, we sought to move beyond correlation and tie these phenotypes directly to bacterial catabolism of the compound. Using *Pantoea sp*. R4, we successfully generated a markerless deletion of *iolG*, which encodes the enzyme that performs the first step in the bacterial catabolic pathway (Figure S3A). This enzyme oxidizes *myo*-inositol to 2-keto-*myo*-inositol and impacts plant colonization in pathogenic and symbiotic microorganisms ^25,28^. While growth curves in minimal media with inositol as a sole carbon source verified this strain lost the ability to catabolize inositol, growth of Δ*iolG* mimicked the parental strain when grown on glucose (Figures 4A and S5A). We posited that loss of this catabolic pathway would result in decreased root colonization.

**Figure 4.**
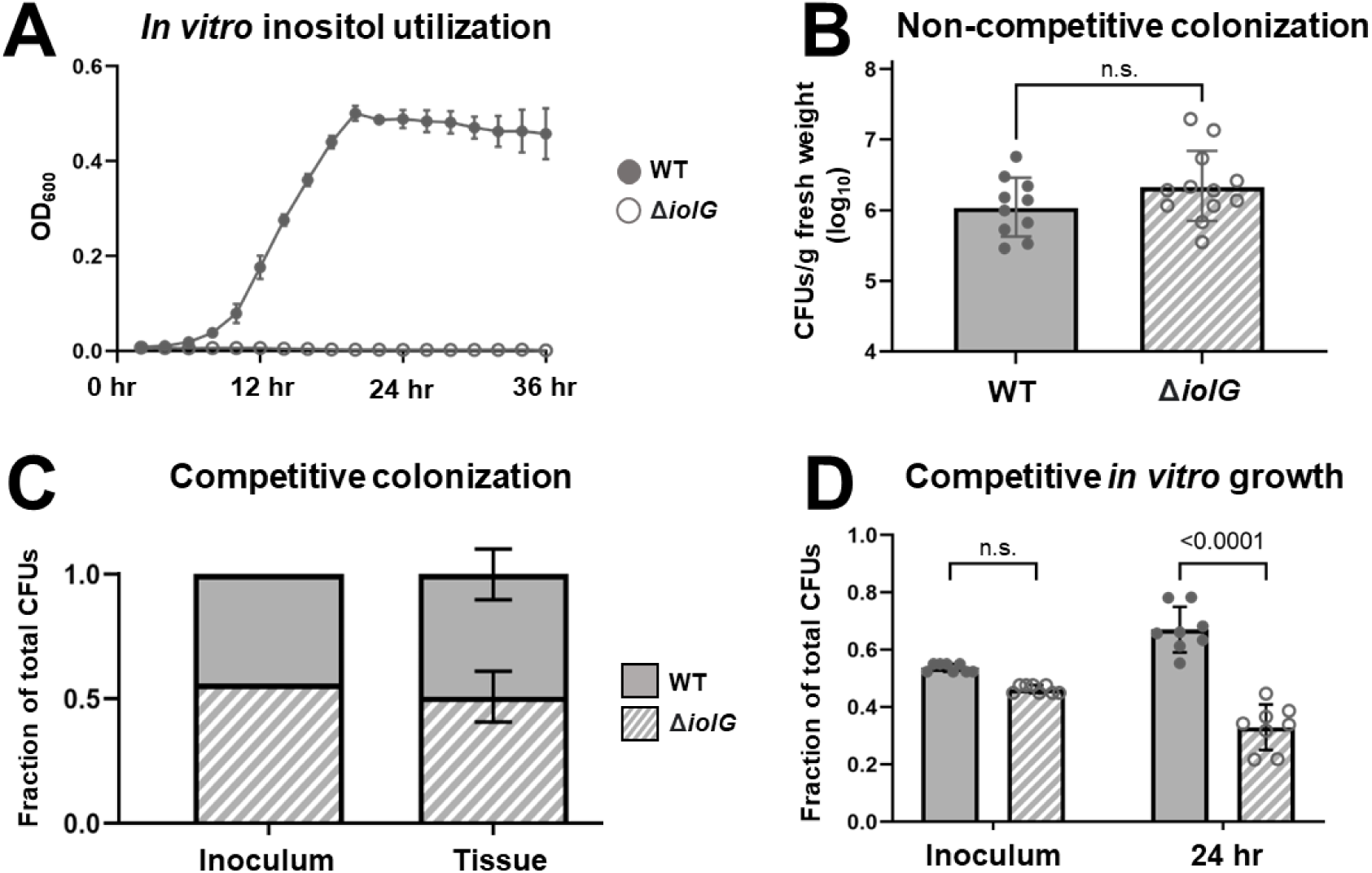
*Pantoea sp*. R4 inositol catabolism does not facilitate increased root colonization but does confer an *in vitro* growth advantage. To causatively examine the effect of inositol catabolism on *Pantoea sp*. R4 root colonization levels, we generated a deletion strain (Δ*iolG*) that cannot catabolize inositol. A) Growth curves of *Pantoea sp*. R4 WT (gray) and Δ*iolG* (white) strains in media with inositol as the sole carbon source. Each curve represents 2 biological replicates with 6 technical replicates each (n=12 wells). Bars represent the standard deviation. B) Colonization of *Pantoea sp*. R4 WT (gray) and Δ*iolG* (hatched) on Arabidopsis Col-0 roots after 1 week. Each symbol represents the pooled roots of 3 seedlings. Data points in panel B fit a lognormal distribution and were therefore log-transformed prior to performing an unpaired t-test with Welch’s correction. Bars represent the geometric mean +/- the geometric SD. C) The two strains were inoculated on Col-0 roots at the ratio shown in ‘Inoculum’. After 7 days, roots were harvested and CFUs from each strain were enumerated. The ratio of each strain in relation to total CFUs was calculated. All replicates received the same inoculum ratio (no error bars), and the ratio after seven days reflected the initial inoculum levels. Data in panel C is from a single biological replicate that included 4 separate growth assays, each with 3 seedlings. A two-way ANOVA (with Šidák’s multiple comparisons test and a single pooled variance) revealed no significant difference between Inoculum and Day 7 ratios, or between WT and Δ*iolG* levels for either the inoculum or tissue samples (p-value < 0.05). Bars represent the mean +/- SD. See also Figure S5E for an additional biological replicate with similar outcomes. D) The WT and Δ*iolG* strain were competitively inoculated into a root homogenate growth substrate. After 24 hours, cells from each strain were enumerated. The ratio of each strain in relation to total CFUs was calculated. Data in panel D is representative of 2 biological replicates, each with 4 technical replicates (n=8 wells). An RM two-way ANOVA (with matched values, Šidák’s multiple comparisons test, and a single pooled variance) revealed no significant difference between WT and Δ*iolG* levels in the inoculum, but a significant difference after 24 hours (p-value < 0.05). Each symbol in panel D represents a single well in a 96-well plate.

When we repeated the colonization assays on Col-0 plants using Δ*iolG*, we failed to observe differences in root colonization levels compared to the parental strain (Figure 4B). Because the effects of a missing enzyme in the first step of the pathway could potentially be rescued if the metabolic byproduct was present in plant tissue during colonization, we generated an additional bacterial mutant strain by deleting 4 enzymes in the inositol catabolic pathway, *iolEGDC*, that are located sequentially in the *Pantoea sp*. R4 genome (Figure S5B). These enzymes constitute 4 of the 5 enzymes in the previously characterized bacterial catabolic pathway (Figure S3A)^28^. We validated the desired phenotype in minimal media with inositol as a sole carbon source (Figure S5C). However, we again observed that when Δ*iolEGDC* was associated with plants for 1 week, there was no difference in colonization levels compared to the parental strain (Figure S5D). Further, we performed co-inoculation assays using the wild-type and Δ*iolG* strains to see if there was a competitive advantage within host tissue. One week after dual inoculation, the recovered CFUs of each strain reflected the initial inoculum levels (Figures 4C and S5E). Altogether, these results indicate that in our experimental set-up, inositol catabolism does not influence *Pantoea sp*. R4 root colonization levels.

### Inositol catabolism provides an in vitro growth advantage

We next moved our experiments outside of the living plant host to test the effects of this catabolic pathway *in vitro*, without the influence of host regulation, compartmentalization, or response to microbial presence. We first homogenized Col-0 seedling roots to create a solution that mimics the nutritional environment of our colonization assays. We then measured the cell density of the *Pantoea* wild-type and Δ*iolG* strain growing in mono-culture or in competition with each other. After 24 hours, there was no difference between the cell densities of each mono-culture compared to the starting inoculum (Figure S5F). On the other hand, co-inoculations revealed the wild-type *Pantoea* strain outcompeted Δ*iolG* (Figure 4D). These assays suggest the inositol catabolic pathway is sufficient to increase short-term growth in a homogenous root nutritional environment and the lack of difference observed in competitive *in vivo* colonization is partially controlled by the living host.

### Inositol sensing increases swimming motility

Motility is an important bacterial behavior during root colonization that can be impacted by the surrounding nutritional environment ^32,33^. We were therefore interested in motility phenotypes between the catabolic mutant and parental *Pantoea* strain. The catabolic mutant is unable to grow when only inositol is supplied, but it still encodes a functional transporter to bring the compound into the cell. When spotted on plates with glucose as a sole carbon source, the wild-type and Δ*iolG* strains display similar levels of motility after 48 hours, as quantified via halo diameter in low-agar media (Figures 5A and 5B). Wild-type *Pantoea sp*. R4 demonstrates increased motility when inositol is the sole carbon source, while the mutant does not grow on the media (Figures 5A). Interestingly, when grown on a combination of glucose and inositol, both strains showed increased motility levels as compared to glucose alone (Figures 5A-5C). It is notable that both the wild-type and catabolic mutant displayed the same increase, even though Δ*iolG* cannot metabolize this compound. This supports a motility phenotype independent of inositol catabolism, but still related to the environmental presence of this compound. We utilized a low-agar media set-up for our plant colonization assays, which would allow for flagellar motility phenotypes to be expressed. These altered behavioral patterns could therefore have contributed to the colonization differences we observed. Together, these results suggest that bacterial behavior is impacted by the presence of inositol in a catabolism-independent manner.

**Figure 5.**
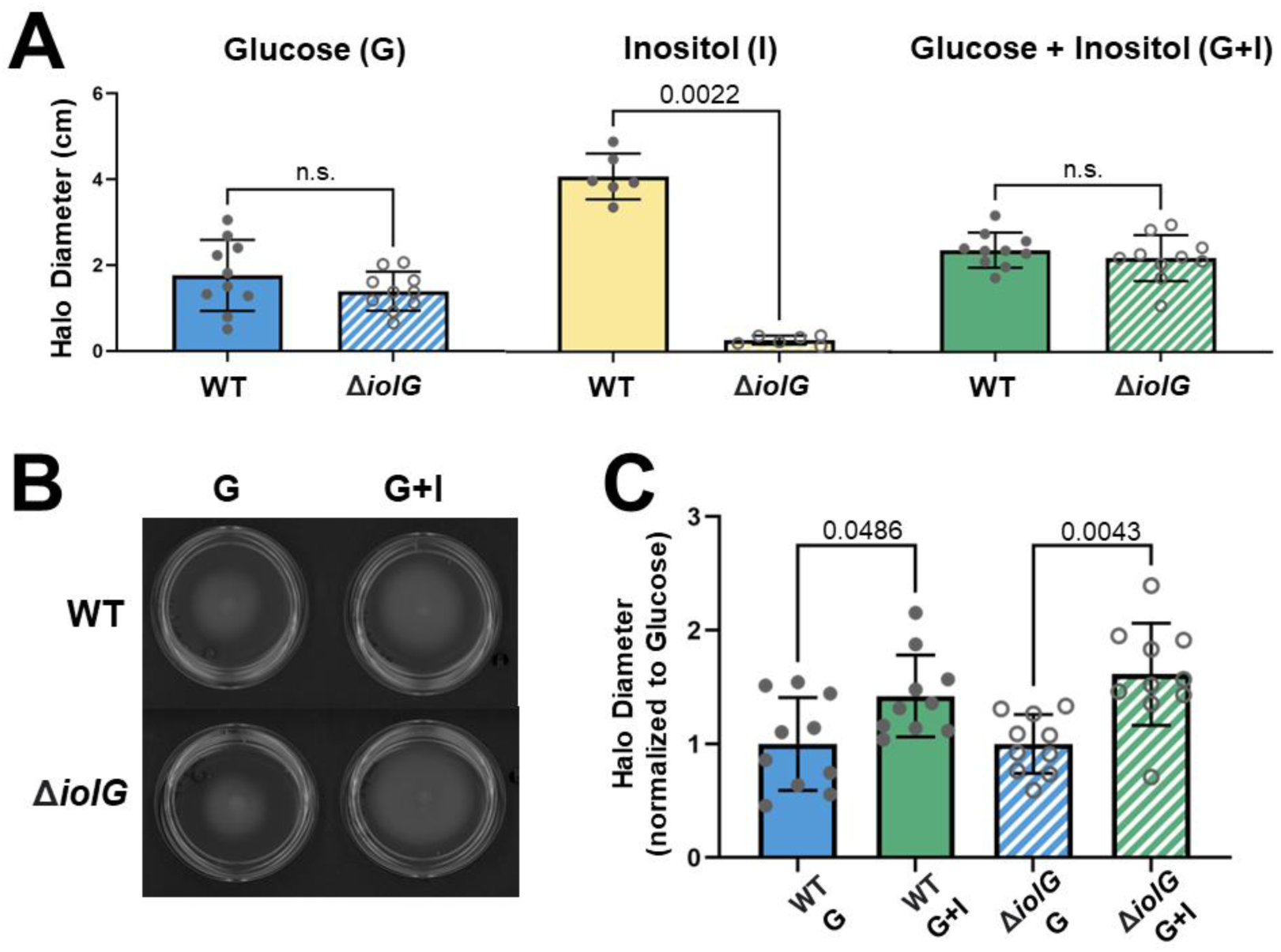
Inositol sensing increases swimming motility. We investigated impacts of bacterial catabolism on a host-relevant phenotype, motility. A) *Pantoea sp.* R4 WT (filled gray circles) and Δ*iolG* (open gray circles) swimming halo diameters were measured on motility plates supplemented with glucose (G, blue shading), inositol (I, yellow shading), and a combination of glucose + inositol (G + I, green shading). Data points for the G and G + I treatments fit a normal distribution and an unpaired t-test with Welch’s correction was performed for each. A non-parametric Mann-Whitney test was performed on data for the I treatment. B) Representative swim plates showcase different halo diameters for *Pantoea sp*. R4 WT and Δ*iolG* on glucose (G) and glucose + inositol (G+I) plates. C) Comparison of *Pantoea sp.* R4 WT (filled gray circles, solid bars) and Δ*iolG* (open gray circles, hatched bars) strain halo diameters grown on G alone (blue) or a combination of G + I (green). Data in panel C was normalized to the respective strain’s halo diameter on glucose. Brown-Forsythe and Welch ANOVA tests (with Dunnett’s T3 multiple comparisons test) were run to compare the nutrient treatments in panel C. Each symbol in panels A and C represents a single plate, and data was collected across at least 2 biological replicates. All bars represent the mean +/- SD. Data in panels A and C are from the same experiments.

## Discussion

Plants grown in distinct environments often harbor similar microbiomes ^31^. This remarkable observation suggests highly conserved mechanisms for the establishment and maintenance of specific plant-microbe interactions. While many host and microbial traits mediating colonization dynamics have been identified, we are far from understanding the full complexity of these relationships. Here, we identify a role for inositol in mediating microbial root colonization phenotypes. Importantly, we provide evidence that host control of this compound mediates plant-microbe interactions across divergent hosts (poplar and Arabidopsis), bacterial phyla (Proteobacteria and Actinomycetes), and abiotic conditions (natural environmental and controlled laboratory conditions).

Inositol is a cyclic polyol used abundantly in eukaryotic cells as a precursor to various structural and signaling molecules ^23^. It can be incorporated into lipid structures as phosphatidylinositol, phosphorylated at multiple locations, and cleaved to release important secondary messengers in response to various signals, such as phytohormones, drought and salt stress, and pathogen perception ^23^. Eukaryotes synthesize inositol through a two-step process that uses glucose-6-phosphate as an initial substrate^34^. First, *myo*-inositol phosphate synthase (MIPS) converts glucose-6-phosphate to inositol-3-phosphate (Fig S2A). An inositol monophosphatase, VTC4, then acts on this compound to produce *myo*-inositol^34^. As a testament to the importance of inositol in various plant functions, Arabidopsis produces three MIPS enzymes that can partially complement each other *in vivo* ^35^. Previous Arabidopsis studies have shown *mips2* and *ipk1* plants have increased susceptibility to bacterial, fungal, and viral pathogens ^36^, but *mips1* plants have increased resistance to an oomycete pathogen ^37^. On the other hand, *vtc4* Arabidopsis plants are impaired in establishing interactions with a beneficial *Bacillus* strain ^24^. While we failed to observe a colonization reduction for our *Pantoea* strain on *mips2*, this could be due to complementation by the other two MIPS enzymes or suggest a differential host response to non-pathogenic microorganisms. Together, these previous works highlight the pleiotropic nature of inositol biosynthesis in mediating diverse plant phenotypes.

We also failed to observe a colonization deficit when we associated *Pantoea* with an Arabidopsis *ipk1* mutant line. This enzyme does not act directly on inositol, but rather phosphorylates downstream inositol phosphates and is involved in the formation of phytic acid, an important and well-studied phosphate storage molecule ^38^. Phytic acid phosphatases have been identified across environments with important ramifications for host interactions ^39^. Inositol phosphate kinases can have a broad range of substrates and there could be additional kinase enzymes that fulfill similar roles to IPK1. While IPK1 did not influence colonization in our experimental set-up, it is important to recognize the crucial role inositol additionally serves as the structural backbone for a major phosphate storage molecule in plants. Inositol can also impact auxin homeostasis and be used for synthesis of the cell wall precursor UDP-glucuronic acid ^40,41^. Therefore, impacts on soluble inositol levels could have a ripple effect on many other aspects of plant physiology, including nutrient, hormone, and cell wall homeostasis. In fact, plant growth is so heavily impacted by inositol that many common plant tissue culture media formulations contain inositol as the sole organic carbon source. This underscores the critical nature of understanding this compound’s role in mediating plant-microbe interactions, as different media usage may unknowingly influence these dynamics.

There are many examples of microbial inositol usage influencing pathogenic and beneficial microbial interactions with diverse plant and mammalian hosts ^24,25,42,43^. For example, altering transport or *de novo* synthesis of inositol reduces the virulence of eukaryotic pathogens that require it for phospholipid synthesis ^43^. While eukaryotes use inositol for many structural and signaling processes, bacteria do not typically lipidate this compound and relatively few bacterial species can synthesize inositol ^44^. Notable exceptions to this are some Actinobacteria, which can incorporate inositol into their membranes or use it to generate redox molecules ^30,44,45^. This is of special interest to our research, as *Streptomyces* are a member of this phylum and the isolate we use encodes a predicted two-step pathway for *de novo* inositol biosynthesis. It is therefore intriguing that we observed comparable colonization reductions between the *Streptomyces* and *Pantoea* isolate, which does not encode this pathway. This highlights a broad role for host-derived inositol in mediating interactions with bacteria that likely have different physiological requirements for this compound.

Bacterial inositol usage is thought to largely center around carbon metabolism. The inositol catabolic pathway is widespread across bacterial genomes, with an impressively high occurrence in rhizosphere-associated organisms and various plant and mammalian pathogens ^2,46^. Various genetics approaches have established a role for microbial inositol catabolism in diverse host-bacteria interactions. For example, the devastating plant pathogen *Ralstonia solanacearum* relies on inositol catabolism during early colonization of tomato roots ^25^, while nitrogen-fixing symbionts require this pathway for competitive host nodulation ^28,47^. Intriguingly, however, our *Pantoea sp.* R4 catabolic mutant did not have a colonization deficit on Col-0 seedlings, but was consistently outcompeted by the parental strain *ex vivo*. Potentially, the heterogeneous environment of the living host, resulting at least in part from transport and compartmentalization of nutrients in response to stimuli, influences colonization outcomes. This idea is further supported by the results of our assays using host genotypes impaired in inositol transport at different cellular locations.

Microbial carbon metabolism is differentially regulated under variable environment conditions ^48^ and influences microbial phenotypes relevant to host interactions, such as biofilm formation and virulence ^49–51^. Importantly, the transcriptional repressor associated with inositol catabolism, *iolR*, has been characterized for several bacteria and influences various aspects of cellular physiology. These include regulating the production of biofilm ^52^ and type III secretion system effectors ^53^, usage of alternative carbon sources ^54^, and reversible lysine acetylation ^55^. Interestingly, the human fungal pathogen *Cryptococcus neoformans* also uses inositol as a sole carbon source ^56^ and this catabolism influences capsule structure, virulence, and other facets of its physiology ^42,57^. Additionally, inositol induces sporulation and mating in diverse fungi, suggesting a broadly conserved role for this compound as an important environmental cue ^58–60^. It is therefore feasible that the observed colonization patterns in our studies are due to the regulatory networks associated with inositol sensing, rather than the catabolic pathway *per se*. Microorganisms that encode this pathway may share conserved regulatory patterns, which would additionally explain our correlative proteobacterial colonization dataset (Figure 3A). The observed *Pantoea* motility phenotypes further support that sensing of this largely eukaryotic-derived compound influences bacterial behavior independently of catabolism. However, motility is only one potential explanatory mechanism for our observed results. While *Pantoea sp.* R4 is motile, *Streptomyces sp*. CL18 is not. Because we observed similar colonization reductions for both of these isolates, it is likely that other altered cellular phenotypes contribute to host interactions.

Intriguingly, the relative importance of inositol via catabolism or the associated regulon may be unique depending on the specific plant-microbe interaction. Previously published supplemental RNA-seq datasets suggest an interesting connection between bacterial expression of this pathway (*iol* pathway) and plant immune status^61–63^. For two well-studied plant pathogens, *Pseudomonas syringae* DC3000 (*Pst* DC3000) and *Ralstonia solanacearum*, *iol* transcripts are decreased *in vivo* compared to rich media, suggesting a conserved response of plant pathogens to downregulate this catabolic pathway during interactions with hosts^61–63^. However, for *Pst* DC3000, *iol* expression significantly increased when the host-pathogen interaction was altered in a variety of ways: 1) activation of host pattern-triggered immunity (PTI) via flg22 pre-treatment before inoculation, 2) colonization of a disarmed, avirulent *Pst* DC3000 (D36E) that cannot suppress PTI, and 3) ectopic expression of the avirulence effector AvrRps4, which activates effector-triggered immunity (ETI)^62^. Together, these patterns suggest that altered *iol* transcripts may be part of the microbial response to the host immune state.

In conclusion, we use a correlative network built on transcriptome profiles of wild poplar trees and their native microbiome to hypothesize a role for host inositol transport in mediating plant-microbe dynamics. We provide causal evidence for these host transporters impacting colonization in a controlled laboratory setting using a different host and individual, phylogenetically distinct microorganisms. Bridging these approaches helps to underscore a robust inositol dependent mechanism influencing interactions across divergent plant and microbial phylogenies, environmental conditions, and community complexities. Intriguingly, we did not observe a direct role for bacterial inositol catabolism in *Pantoea* colonization outcomes, despite robust evidence for this pathway’s importance in varied pathogenic and beneficial interactions. We did, however, observe effects of inositol on microbial behavior that were independent of catabolism, suggesting inositol sensing alters other aspects of microbial physiology. Although we only verified a motility phenotype mediated by this compound, we cannot rule out many other host-relevant phenotypic alterations that may occur in response to this common root exudate, several of which have been observed in other plant and mammalian host-microbe systems. Our data, together with others, suggests host control of this compound and resulting microbial behavior are important mechanisms at play surrounding the host metabolite inositol. However, further work is needed to fully understand the mechanistic underpinnings of this host-microbe-metabolite interaction network.

## Acknowledgements

Several isolates used in this study were originally isolated in the laboratory of Jeffery Dangl at the University of North Carolina. We would like to thank the labs of Dr. Erik Zinser and Dr. Gladys Alexandre, as well as Elise Phillips, Trevor Hancock, and Alexandra Gates for generously providing important strains, reagents, and discussion for this work. This work is supported by the National Science Foundation grants IOS-1750717 to S. Lebeis, DEB-1638922 to S. Lebeis, and DGE-1938092 to B. O’Banion. Any opinions, findings, and conclusions or recommendations expressed in this material are those of the author(s) and do not necessarily reflect the views of the National Science Foundation. This work was additionally supported by the Science Alliance Joint Directed Research and Development Funding at Oak Ridge National Laboratory (awarded to S. Lebeis and D. Jacobson) and the Tennessee Plant Research Center (awarded to B. O’Banion). This research used resources of the Oak Ridge Leadership Computing Facility, which is a DOE Office of Science User Facility supported under Contract DE-AC05-00OR22725. Funding was provided by the Plant-Microbe Interfaces (PMI) and by The Center for Bioenergy Innovation (CBI), these are both supported by the Genomic Sciences Program of the Office of Biological and Environmental Research in the DOE Office of Science. The metatranscriptome sequencing conducted by the US Department of Energy Joint Genome Institute is supported by the Office of Science of the US Department of Energy under Contract no. DE-AC02-05CH11231. Support for the Poplar GWAS SNP dataset is provided by the U.S. Department of Energy, Office of Science Biological and Environmental Research (BER) via the Bioenergy Science Center (BESC) under Contract No. DE-PS02-06ER64304. The Poplar GWAS Project used resources of the Oak Ridge Leadership Computing Facility and the Compute and Data Environment for Science at Oak Ridge National Laboratory, which is supported by the Office of Science of the U.S. Department of Energy under Contract No. DE-AC05-00OR22725. This manuscript has been co-authored by UT-Battelle, LLC under contract no. DE-AC05-00OR22725 with the U.S. Department of Energy. The United States Government retains and the publisher, by accepting the article for publication, acknowledges that the United States Government retains a nonexclusive, paid-up, irrevocable, world-wide license to publish or reproduce the published form of this manuscript, or allow others to do so, for United States Government purposes. The Department of Energy will provide public access to these results of federally sponsored research in accordance with the DOE Public Access Plan (http://energy.gov/downloads/doe-public-access-plan, last accessed September 16, 2020).

## Author Contributions

Conceptualization, B.S.O, D.J., and S.L.L.; Methodology, B.S.O, P.C.J., D.J., S.L.L.; Software, P.C.J. and D.J.; Formal Analysis, B.S.O. and P.C.J.; Investigation, B.S.O., P.C.J., A.A.D., A.S.W., and B.R.K.; Resources,J.C., W.M., T.B.R., D.J., and S.L.L.; Data Curation, B.S.O. and P.C.J.; Writing - Original Draft, B.S.O and S.L.L.; Writing - Review and Editing, B.S.O., P.C.J., A.A.D., A.S.W., B.R.K., T.B.R., D.J., and S.L.L.; Visualization, B.S.O.; Supervision, T.B.R., D.J., and S.L.L.; Funding Acquisition, B.S.O., D.J., and S.L.L.

## Methods

### GWAS dataset generation and sub-network curation

The GWAS was performed according to the detailed methods in Weighill, et al^64^. Briefly, we obtained total RNA-seq data for leaf and xylem tissues of *Populus trichocarpa* genotypes and aligned trimmed and filtered reads to the *P. trichocarpa* v3.0 reference genome from Phytozome ^21,65,66^. Unmapped reads were assumed to be putative members of the microbiome and used for metatranscriptome analysis. These unmapped reads were assigned to a taxa using ParaKraken and aggregated at the genus level ^67^. We filtered for taxa that were present with a relative pseudo-abundance of at least 1%, performed sample level normalization to mitigate sequencing bias, and discarded putative outlier taxa. The remaining microbial taxa were used as phenotypes against poplar SNPs, after applying a minor allele frequency cutoff of 0.05. Significant SNP to phenotype association was determined using a linear mixed model approach implemented in EMMAX^68^, applying a two-stage FDR threshold. We visualized these taxa to gene associations in a Cytoscape network and used MapMan ontologies for functional annotation ^22,69^. Specifically, we focused on reads that mapped to the genera *Pantoea* and *Streptomyces*.

### Plant lines and maintenance

To identify Arabidopsis mutants of interest, poplar GWAS results were aligned to the Arabidopsis genome (TAIR10) using Phytozome^66^. The best Arabidopsis match (via lowest e-value) was used as a gene query for finding T-DNA insertion line seed stocks in The Arabidopsis Information Resource online portal^70^. The selected lines were then obtained from the Arabidopsis Biological Resource Center (ABRC, Ohio State University) ^71^. The obtained seed stocks are as follows: *int1* (AT2G43330, SALK_123915C); *pmt5* (AT3G18830, SALK_005175C); *mips2* (AT2G22240, SALK_031685C), and *ipk1* (AT5G42810, SALK_075528C). All germplasms are in a Col-0 background. All seeds were surface sterilized in 70% ethanol with 0.1% Triton X-100 for 3 min, 10% household bleach with 0.1% Triton X-100 for 15 min, followed by three washes with sterile distilled water. Seeds were stratified for at least 3 days in the dark at 4°C and subsequently germinated at 24°C with 16 h of light for 8 to 11 days on agar plates containing half-strength (2.22 g/L) Murashige & Skoog (MS) basal medium, 1% sucrose, and 0.8% Phytoagar (Bioworld). MS basal medium with Gamborg’s vitamins (with *myo*-inositol, base catalog #2623220 from MP Biomedical) was used for germination plates in the 2-week Proteobacterial magenta jar colonization assays (Figures 3A-3C). In all other experiments, MS basal salt mixture (without *myo*-inositol, base catalog #2623022 from MP Biomedical) was used for germination plates.

### Bacterial isolates and inoculum preparation

We used a collection of bacterial strains isolated from a variety of plants and soils (Table S1). Bacterial isolates were grown in lysogeny broth (LB)^72^ at 30°C with shaking at 150 rpm for 1 to 3 days. Cultures were pelleted via centrifugation, resuspended in an equivalent volume of 1x PBS, and normalized to an OD_600_ of 0.01 using 1x PBS. Inoculum of *Streptomyces sp*. CL18 was prepared slightly differently because it forms aggregates in liquid media. Therefore, after growth in LB as described above for 3-5 days, aggregates were placed in a microcentrifuge tube with 1x PBS and 3mm glass beads. The culture was homogenized using a 2010 Geno/Grinder at 1,400-1,500 rpm for 5 minutes, and then normalized to an OD_600_ of 0.1 for further work.

### Bacterial colonization experiments

For 2-week Proteobacterial magenta jar colonization assays (Figures 3A-3C), jars were filled with 100 mL of quarter-strength (1.11 g/L) MS basal medium with Gamborg’s vitamins (with *myo*-inositol, base catalog #2623220 from MP Biomedical) containing 0.3% Phytoagar. Five Arabidopsis Col-0 seedlings were aseptically placed equi-distant from each other, and 150 μL of bacterial isolate resuspensions at OD_600_ 0.01 were injected at a distal location. Jars were sealed with BreatheEasy film and placed in a growth chamber at 24°C with 16 h of light for 14 days. For all other colonization experiments in 12-well plates, quarter-strength (1.11 g/L) MS basal salt mixture (without *myo*-inositol, base catalog #2623022 from MP Biomedical) containing 0.3% Phytoagar was pre-mixed with bacterial inoculum (1.5 μL of OD_600_ 0.01 resuspensions/mL [OD_600_ 0.1 for *Streptomyces* assays]) and dispensed into individual wells (3 mL/well). For the dual-inoculation treatments, equal volumes of each normalized culture were mixed prior to adding to the MS media. Three Arabidopsis seedlings were aseptically transferred to each well using flame-sterilized tweezers. Tissue-culture plates were sealed with Parafilm M Laboratory Film and placed in a growth chamber at 24°C with 16 h of light for 7 days. In treatments where exogenous inositol was added, MS basal medium with Gamborg’s vitamins (with *myo*-inositol, base catalog #2623220 from MP Biomedical) was used instead. In addition to *myo*-inositol (100 mg/L), the MS basal medium with Gamborg’s vitamins also contains an addition of 1 mg/L nicotinic acid, 1 mg/L pyridoxine-HCl, and 10 mg/L thiamine-HCl.

After the allotted time frame for each experimental set-up (1 or 2 weeks), seedlings were aseptically harvested using flame-sterilized tweezers and scissors to separate leaf and root tissue into sterile, pre-weighed 1.5-mL centrifuge tubes. After seedling removal, the surrounding MS media was mixed and 1 mL was removed for quantification of bacterial survival in the bulk media. Tissue from the same jar (5 seedlings) or well (3 seedlings) was pooled together for downstream analyses. Tubes were weighed again after tissue addition to determine fresh-weight biomass. One mL of 1x PBS was added to root tissue samples and vortexed vigorously for 30 seconds to remove loosely-attached microorganisms. Roots were then removed and placed into new microcentrifuge tubes. The remaining 1 mL 1x PBS was retained for rhizosphere colonization enumeration. The transferred root tissue was subsequently washed twice with sterile, distilled H_2_O. We added 3mm glass beads to the microcentrifuge tube containing root tissue and homogenized the samples using a 2010 Geno/Grinder at 1,400-1,500 rpm for 5-10 min. The root homogenate was diluted and plated on LB plates to enumerate internal and tightly attached external bacteria. For competitive colonization assays, samples were plated on 1x M9 minimal salts with 1 mM MgSO_4_ and 10 mM of *myo*-inositol. The WT *Pantoea* strain forms robust colonies on M9 + inositol, while the *ΔiolG* strain forms very small dots, likely due to residual carry-over from other compounds in the growth assay. This difference in colony morphology facilitated counting both strains on a single plate.

### Phylogenetic tree construction

All bacterial isolates used in this study were previously sequenced and are available in public repositories (Table S1). Whole genomes for each isolate were downloaded from the Joint Genome Institute’s IMG/MER database and run through AMPHORA to scan for 31 house-keeping protein-coding genes ^73,74^. We removed 4 genes from further analysis because they were not found in a single copy across all genomes. The remaining 27 genes were aligned using MAFFT (version 7) ^75^ on the CIPRES gateway ^76^ and concatenated in MEGA (version 11.0.11) ^77^. The concatenated alignments were used to build a maximum likelihood phylogenetic tree using RAxML (version 8.2.12)^78^ with 1000 bootstraps on the CIPRES gateway. Tree visualization was done in iTOL and graphical editing done in Adobe Illustrator ^79^.

### Bacterial growth curves

Bacterial isolates were grown in LB broth as described above. Cells were washed to remove excess nutrients by centrifuging 1 mL of turbid culture at 13,300 g for 3 minutes, decanting supernatant, resuspending in 1x M9 Minimal Salts + 1 mM MgSO_4_, and repeating this wash three times. Resuspensions were normalized to an OD_600_ of 0.01 using 1x M9 minimal salts with 1 mM MgSO_4_ and 10 mM of either *myo*-inositol, glucose, or malate. Resuspended cultures were then dispensed into clear 96-well plates (200 μL/well), incubated at 30°C while shaking, and OD_600_ measurements were taken continuously for 3 days. Blank well readings were subtracted from all growth measurements to remove background readings. For calculations in Table S2, the average OD_600_ of each isolate growing on inositol after 72 hours was divided by the average OD_600_ for the reference carbon source (glucose or malate). Growth data was visualized in Graphpad Prism.

### Plasmid construction

For all work, plasmid maintenance was performed using *E. coli* Dh5α. The *E. coli* Dh5α strain harboring an empty pk18mobsacB vector (gift of G. Alexandre) was grown in LB + Kanamycin (Kan, 30 μg/mL) and plasmid extractions were done using the Wizard Plus SV Minipreps DNA Purification System according to the manufacturer’s protocol (Promega). The pk18mobsacB vector contains a kanamycin resistance cassette and the *sacB* gene, which confers sensitivity to sucrose^80^. All primers, plasmids, and target genes are further described in Table S4.

To generate the *iolG* deletion cassette, approximately 1000bp of the 5’ and 3’ untranslated region (UTR) flanking *iolG* was amplified with primers BSO1/BSO2 and BSO3/BSO4, respectively. These primer sets introduced a 30bp complementary overhang to the 3’ end of the 5’ UTR PCR product and the 5’ end of the 3’ UTR PCR product. This complementarity was used to splice the fragments together using overlap extension PCR (OE-PCR). Following splicing, BSO1 and BSO4 were used for further amplification of the OE-PCR product. After successful amplification of the stitched 5’ and 3’ UTR regions, a SalI cut site was introduced to the 5’ and 3’ end of the stitched PCR product with primers BSO5/BSO6. The resulting PCR product and the pk18mobsacB empty vector were both digested with SalI and ligated to generate pk18mobSacB-iolG.

To generate the *iolEGDC* deletion cassette, approximately 1000bp of the 5’ UTR flanking the *iolEGDC* genomic region was amplified with primers BSO7/BSO8, which introduced a XmaI and HindIII cut site to the 5’ and 3’ end of the PCR product, respectively. Additionally, approximately 1000bp of the 3’ UTR flanking the *iolEGDC* genomic region was amplified with primers BSO9/BSO10, which introduced a HindIII and NheI cut site to the 5’ and 3’ end of the PCR product, respectively. The 3’ UTR PCR product and the pk18mobsacB vector were digested with HindIII and NheI and ligated together. Next, the 5’ UTR PCR product and the pk18mobsacB vector containing the 3’ region were digested with XmaI and HindIII and ligated together, creating pk18mobsacB-iolEGDC.

For both pk18mobsacB-iolG and pk18mobsacB-iolEGDC, the ligated constructs were heat-shocked into NEB5α competent *E. coli* cells and individual colonies were genetically screened for the correct construct.

### Generation of *Pantoea sp*. R4 deletion strains

*E. coli* strains containing the plasmids of interest were grown to mid-log phase for plasmid extractions. The isolated plasmid was then heat-shocked into *E. coli* EZ180 competent cells (gift of E. Zinser). EZ180 is a diaminopimelic acid (DAP) auxotroph. A lawn of EZ180 (LB + Kan + DAP, 37°C) and *Pantoea sp*. R4 (LB, 30°C) were each grown overnight. The resulting lawns were removed and re-plated on LB + DAP in approximately equal densities and left to conjugate at 30°C overnight. The mixed growth lawn was removed by adding 1 mL LB broth to the plate, resuspending the lawn growth, and collecting it in a microcentrifuge tube. This mixture was plated on LB + Kan to select for *Pantoea sp*. R4 cells that went through homologous recombination with the plasmid DNA. Once single recombinants were identified, they were grown overnight in LB broth without antibiotic selection 3 times. Serial dilutions were then plated on 1x M9 Minimal Salt plates containing 10% sucrose (w/v) to select for double recombination events. Colonies that appeared on plates containing 10% sucrose were rescreened for growth on LB and loss of growth on LB + Kan, suggesting the cassette had been kicked out of the genome. Putative colonies were then genetically screened for successful markerless deletion and phenotyped in 1x M9 Minimal Salt medium containing inositol as the sole carbon source.

### Root homogenate growth assays

Arabidopsis Col-0 seeds were sterilized, vernalized, and germinated as described above. After 10 days, the roots of approximately 20 seedlings were placed in a 5 mL tube with 2 mL of quarter-strength MS basal salt mixture (without *myo*-inositol, base catalog #2623022 from MP Biomedical) and 3 mm glass beads. Roots were homogenized using a 2010 Geno/Grinder at 1,400-1,500 rpm for ∼8 min. The resulting homogenate was filter-sterilized with a 0.2 μm filter and stored at 4°C until use. To set-up the growth assays, a homogenous root mixture was vortexed and aliquoted into 1.5 mL centrifuge tubes to ensure each subsequent inoculum contained an identical starting nutrient regime. Overnight bacterial cultures were centrifuged, washed, and added to the root mixtures for a final OD_600_ of 0.01. For the dual-inoculation treatment, equal volumes of each normalized culture were mixed prior to adding to the root mixture. Next, 100 μL of each treatment was inoculated in biological quadruplicate into wells of a clear, flat-bottomed 96-well plate and incubated statically at 30°C. A portion of the inoculum was immediately plated for enumeration. After 24 hours, each well was mixed using a pipette and a 10 μL aliquot was removed for serial dilution plating on 1x M9 minimal salts with 1 mM MgSO_4_ and 10 mM *myo*-inositol as described above.

### Genome Comparisons

We compared the genomes of the 12 Proteobacterial isolates using the Phylogenetic profiler for single genes through the Joint Genome Institute’s IMG/MER toolkit (max e-value = 1^10^-5; minimum percent identity = 30%; algorithm = presence/absence; minimum taxon percent with homologs = 100%; minimum taxon percent without homologs = 100%) ^81^. Additionally, we used Anvi’o for a complementary genomic comparison ^82^. All genomes were downloaded from JGI IMG/MER (Fasta format, nucleic acids). Genes were annotated with NCBI COGs using DIAMOND (--sensitive flag) ^83^ and a genomes storage database was created for all 12 Proteobacterial isolates. Pan-genomes were built using NCBI blastp ^84^ and one of three MCL inflation thresholds (1, 2, or 5) ^85^. In both analyses, we asked which genes (or gene clusters) were present in inositol ‘users’ (8 genomes), but absent from ‘non-users’ (4 genomes). The gene outputs are described in Table S3. Due to sequence divergence among our isolates, some of the *iol* genes were placed in multiple distinct clusters and therefore were not revealed by our methods.

### Motility Assays

Bacterial cultures were prepared as described above. Soft agar motility plates were prepared using 1x M9 Minimal Salts with 1 mM MgSO_4_, 10 mM of either *myo*-inositol or glucose, and 0.3% agar. For plates with a combination of glucose and inositol, 5 mM of each was added (10 mM total carbon). For motility assays, 5 μl of a washed and normalized (OD_600_ of 0.1) bacterial culture was spotted in the middle of each plate and incubated statically at 30°C. Motility halos were scanned and measured after 48 hours of incubation.

### Statistical analysis

All statistics were performed using GraphPad Prism version 9.4.1 for Windows, GraphPad Software, San Diego, California USA, www.graphpad.com. All datasets were first tested for normality and lognormality using the D’Agostino & Pearson test. If data fit a normal distribution, parametric statistical tests were performed on the raw data. If data was non-normal but fit a lognormal distribution, the raw data was log-transformed prior to performing parametric tests. If neither test was passed, non-parametric tests were used. Specific tests are indicated in each figure legend.

### Resource Availability

#### Lead Contact

Further information and requests for resources and reagents should be directed to and will be fulfilled by the Lead Contact, Sarah Lebeis.

#### Materials Availability

Plasmids generated in this study will be made available on request, but we may require a payment and/or a completed Materials Transfer Agreement if there is potential for commercial application.

#### Data and Code Availability

The GWAS was performed according to the detailed methods in Weighill, et al^64^ and was built using the RNA-seq reads described in Zhang, et al ^21^. Further information on data deposition and algorithms used can be found in these manuscripts. Further inquiries can be directed to the corresponding authors.

**Dataset S1.** Poplar genes identified by GWAS sub-network analysis.

