## Supplemental Information for "Plant inositol transport influences bacterial colonization phenotypes"

### Supplementary Tables

| **Wild-type Strains** | | | | | | |
|---|---|---|---|---|---|---|
| Full Name | Shortened Name | JGI IMG Genome ID | Family | Isolation Source | Encodes pathway? | User? |
| *Pantoea* sp. AlfR4 | *Pantoea* sp. R4 | 2824860516 | Erwiniaceae | Alfalfa | Yes | Yes |
| *Streptomyces* sp. UNC401CLCol | *Streptomyces* sp. CL18 | 2563366515 | Streptomycetaceae | Arabidopsis | Yes | Yes* |
| *Rhizobium* sp. 2MFCol3.1 | *Rhizobium* 2 | 2517572231 | Rhizobiaceae | Arabidopsis | Yes | Yes |
| *Agrobacterium* sp. 33MFTa1.1 | *Agrobacterium* 33 | 2561511224 | Rhizobiaceae | Arabidopsis | Yes | Yes |
| *Ochrobactrum* sp. 370MFChir3.1 | *Ochrobactrum* 370 | 2643221500 | Brucellaceae | Arabidopsis | Yes | Yes |
| *Azospirillum brasilense* sp7 | *Azospirillum* sp7 | 2597490356 | Azospirillaceae | Wheat | No | No |
| *Variovorax paradoxus* CL14 | *Variovorax* CL14 | 2643221508 | Comamonadaceae | Arabidopsis | No | No |
| *Janthinobacterium* sp. TND4EL3 | *Janthinobacterium* EL3 | 2708742400 | Oxalobacteraceae | Duckweed | Yes | Yes |
| *Ralstonia* sp. UNC404CL21Col | *Ralstonia* CL21 | 2558309150 | Burkholderiaceae | Arabidopsis | No | No |
| *Burkholderia* CL11 | *Burkholderia* CL11 | 2546825541 | Burkholderiaceae | Arabidopsis | Yes | Yes |
| *Dyella japonica* UNC79MFTsu3.2 | *Dyella* 79 | 2556921674 | Rhodanobacteraceae | Arabidopsis | No | No |
| *Pseudomonas* sp. TN43 | *Pseudomonas* TN43 | 2824522127 | Pseudomonadaceae | Soil | Yes | Yes |
| *Enterobacter* sp. Sphag71 | *Enterobacter* 71 | 2824531566 | Enterobacteriaceae | Sphagnum moss | Yes | Yes |
| **Deletion Strains** | | | | | | |
| Parental Strain Name | Deletion Strain Name | | | | | |
| *Pantoea* sp. R4 | *Pantoea* sp. R4 Δ*iolG* | | | | | No |
| *Pantoea* sp. R4 | *Pantoea* sp. R4 Δ*iolEGDC* | | | | | No |

**Table S1.** List of wild-type and deletion bacterial strains used in this study. **Streptomyces* sp. CL18 can use inositol as a sole carbon source but was not included as a “user” in the comparative genomic analyses.

| **Strain** | **Glucose (reference)** | **Malate (reference)*** | **Relative Inositol Growth** | **User/Non-User** |
|---|---|---|---|---|
| *Pantoea* sp. R4 | 100% | **-** | 89.51% | User |
| *Rhizobium* 2 | **-** | 100% | 98.24% | User |
| *Agrobacterium* 33 | 100% | **-** | 130.41% | User |
| *Ochrobactrum* 370 | 100% | **-** | 152.13% | User |
| *Azospirillum* sp7 | **-** | 100% | 1.60% | Non-User |
| *Variovorax* CL14 | 100% | **-** | 0.07% | Non-User |
| *Janthinobacterium* EL3 | ** | ** | ** | User |
| *Ralstonia* CL21 | 100% | **-** | 0.65% | Non-User |
| *Burkholderia* CL11 | 100% | **-** | 69.32% | User |
| *Dyella* 79 | 100% | **-** | 1.59% | Non-User |
| *Pseudomonas* TN43 | 100% | **-** | 112.59% | User |
| *Enterobacter* 71 | 100% | **-** | 85.27% | User |

**Table S2.** The ability of each strain to grow on inositol (as measured by $OD_{600}$) as compared to a reference carbon source was calculated. The average $OD_{600}$ at 72 hours post-inoculation was averaged across all replicates for each isolate. The $OD_{600}$ when grown on inositol was divided by the reference carbon source (either glucose or malate), to determine the relative inositol growth. *Malate was used as a positive control carbon source for organisms that are unable to utilize glucose. **EL3 forms aggregates in broth media, so accurate OD readings were unattainable.

| Approach | Gene Name | IMG gene ID (JGI IMG/MER only) | Clustering Parameter (Anvi'o only) | *iol* pathway gene |
|---|---|---|---|---|
| JGI IMG/MER | SulP family sulfate permease | 2824862778 | N/A | N/A |
| | 3D-(3,5/4)-trihydroxycyclohexane-1,2-dione acylhydrolase | 2824863954 | N/A | *iolD* |
| | 5-dehydro-2-deoxygluconokinase | 2824863955 | N/A | *iolC* |
| | 5-deoxy-glucuronate isomerase | 2824863958 | N/A | *iolB* |
| Anvi'o | 5-deoxy-D-glucuronate isomerase | N/A | MCL 1 | *iolB* |
| | Multidrug transporter EmrE and related cation transporters | N/A | MCL 1 | N/A |
| | 5-deoxy-D-glucuronate isomerase | N/A | MCL 2 | *iolB* |
| | 5-deoxy-D-glucuronate isomerase | N/A | MCL 5 | *iolB* |

**Table S3.** Genes identified by comparative genomics approaches. The IMG Gene ID's reference the appropriate gene in *Pantoea sp*. R4's genome. Anvi'o was run with 3 different MCL clustering parameters (1, 2, or 5).

| | Name | Sequence (5' to 3') | Description | Source |
|---|---|---|---|---|
| **Primers** | BSO_01 | GACATGATTACGAATTCGAGCTCGGTACCCGCTGAAGTAATAAAAAAGGGCGGCATTATG | amplify 1kb 5' UTR of *iolG* (fwd); used in SOE PCR of 5' and 3' UTR fragments | This Study |
| | BSO_02 | GATAACCCTCTGAACCCTGTAATCTGGAGTCATTTTTCTCTCTCTGGACGCTTGCGTCTTC | amplify 1kb 5' UTR of *iolG* (rev) | This Study |
| | BSO_03 | GAAGACGCAAGCGTCCAGAGAGAGAAAAATGACTCCAGATTACAGGGTTCAGAGGGTTATC | amplify 1kb 3' UTR of *iolG* (fwd) | This Study |
| | BSO_04 | CATGCCTGCAGGTCGACTCTAGAGGATCCCCAAGCTGGATGGCGTGCAGCTGCTGGCTGATG | amplify 1kb 3' UTR of *iolG* (rev); used in SOE PCR of 5' and 3' UTR fragments | This Study |
| | BSO_05 | AACTTGTCGACGACATGATTACG | used to introduce SalI cut site to 5' of SOE PCR product | This Study |
| | BSO_06 | CATGCCTGCAGGTCGAC | used to introduce SalI cut site to 3' of SOE PCR product | This Study |
| | BSO_07 | AACTTCCCGGGGCAGTGGTGGGTTGCT | amplify 1kb 5' UTR of *iolEGDC* genomic region (fwd); introduced XmaI cut site | This Study |
| | BSO_08 | AACTTAAGCTTAAACCGCCGGGATCTAATC | amplify 1kb 5' UTR of *iolEGDC* genomic region (rev); introduced HindIII cut site | This Study |
| | BSO_09 | AACTTAAGCTTCGGGCGGCCTTTTTAAG | amplify 1kb 3' UTR of *iolEGDC* genomic region (fwd); introduced HindIII cut site | This Study |
| | BSO_10 | AACTTGCTAGCGCTATCCGGCAAACACAAC | amplify 1kb 3' UTR of *iolEGDC* genomic region (rev); introduced NheI cut site | This Study |
| **Plasmids** | pk18mobSacB | N/A | Empty vector; $Suc^s$ (*sacB*), $Kan^R$ | Schäfer et al. 1994 |
| | pk18mobSacb-iolG | N/A | Vector used to generate unmarked *Pantoea sp*. R4 Δ*iolG* mutant | This Study |
| | pk18mobsacB-iolEGDC | N/A | Vector used to generate unmarked *Pantoea sp*. R4 Δ*iolEGDC* mutant | This Study |

**Table S4.** List of primers and plasmids used for cloning in this study.

#### Supplementary Figures

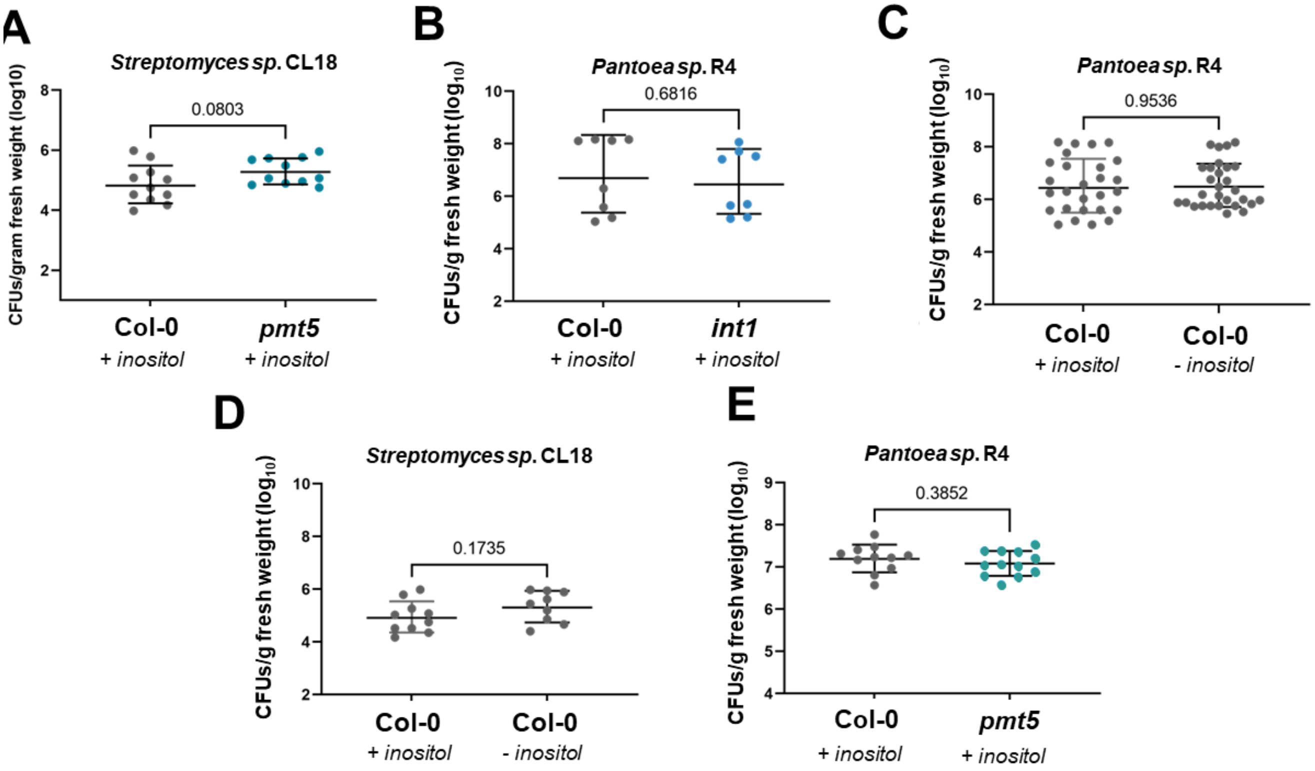


**Figure S1. Exogenous inositol rescues the colonization deficit on *int1* and *pmt5* plants, but does not alter colonization levels on Col-0 plants.** To investigate the role of exogenous inositol in colonization phenotypes, we performed 7-day colonization assays using representative bacterial isolates and Arabidopsis T-DNA insertion lines (*pmt5* and *int1*). A) *Streptomyces sp*. CL18 (square symbols) root colonization levels on Col-0 and *pmt5* seedlings in the presence of exogenous inositol. B) *Pantoea sp*. R4 (circular symbols) root colonization levels on Col-0 and *int1* seedlings in the presence of exogenous inositol. The root colonization of *Pantoea sp*. R4 (C) and *Streptomyces sp*. CL18 (D) on Col-0 seedlings with and without exogenous inositol. E) *Pantoea sp*. R4 root colonization levels on Col-0 and *pmt5* seedlings in the presence of exogenous inositol. Each symbol represents the pooled roots of 3 seedlings (gray dots = Col-0, teal dots = *pmt5*, and blue dots = *int1*). Data points from all panels fit a lognormal distribution and were therefore log-transformed prior to performing an unpaired t-test with Welch's correction. Actual p-values are displayed on each graph. Bars represent the geometric mean +/- geometric standard deviation (SD). Inositol treatment (+/- inositol) is labeled in all panels.

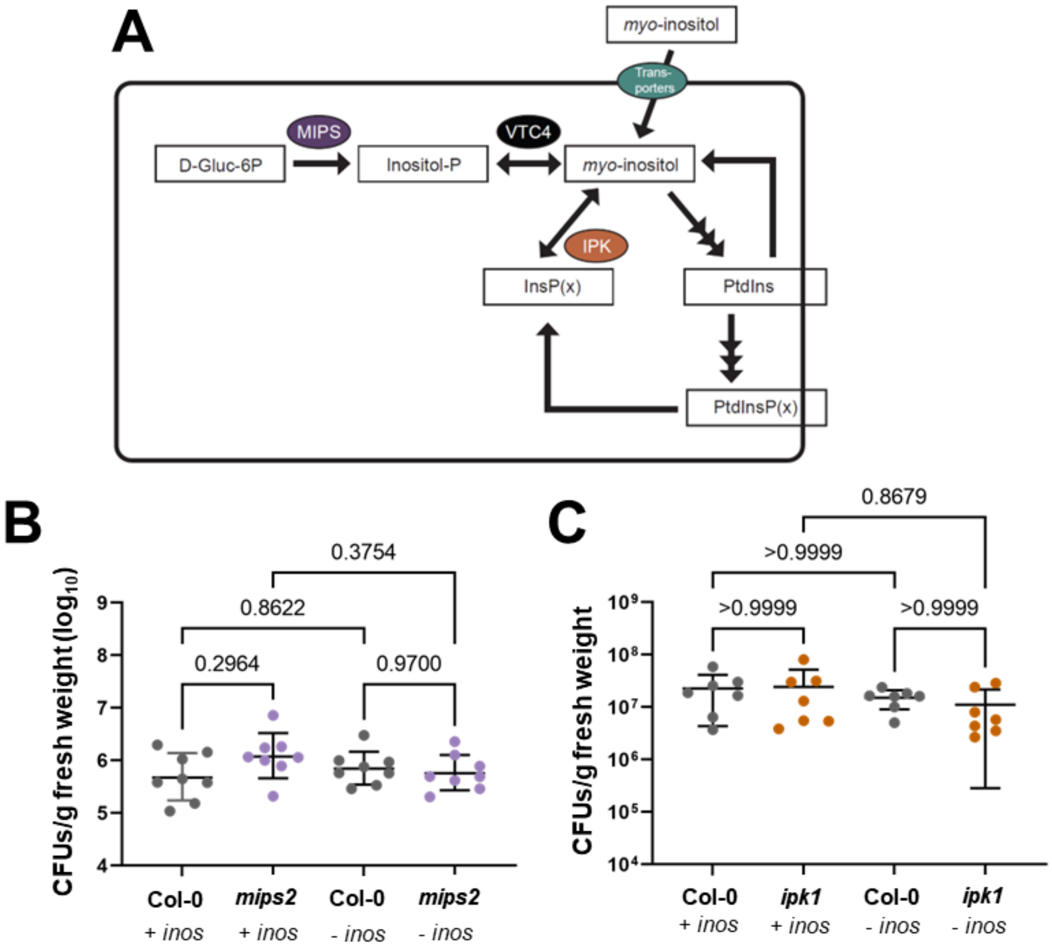


**Figure S2. Mutations in inositol biosynthesis and downstream transformation do not influence *Pantoea sp.* R4 colonization.** A) A simplified diagram showing relevant steps in eukaryotic transport, biosynthesis, and transformation of inositol. Squares represent chemical compounds and ellipses represent proteins or enzymes. B) *Pantoea sp*. R4 root colonization levels between Col-0 and *mips2* seedlings with and without exogenous inositol. C) *Pantoea sp*. R4 root colonization levels between Col-0 and *ipk1* seedlings with and without the presence of exogenous inositol. Each symbol represents the pooled roots of 3 seedlings (gray dots = Col-0, purple = *mips2*, and orange = *ipk1* [colors mimic enzyme symbols in panel A]). Data points in panel B fit a lognormal distribution and were therefore log-transformed prior to performing Brown-Forsythe and Welch ANOVA tests (with a Dunnett's multiple comparisons test). Bars in panel B represent the geometric mean +/- geometric SD. A non-parametric Kruskal-Wallis test (with a Dunn's multiple comparisons test) was performed on data in panel C, where the bars represent the mean +/- SD. Actual p-values are displayed on each graph. Inos = inositol; D-Gluc-6P= D-Glucose-6-Phosphate; Inositol-P = inositol-3-phosphate; InsP(x) = various phosphorylated inositol phosphates; PtdIns = phosphatidylinositol; PtdInsP(x) = various phosphorylated phosphatidylinositol moieties. Inositol treatment (+/- inos) is indicated in panels B and C.

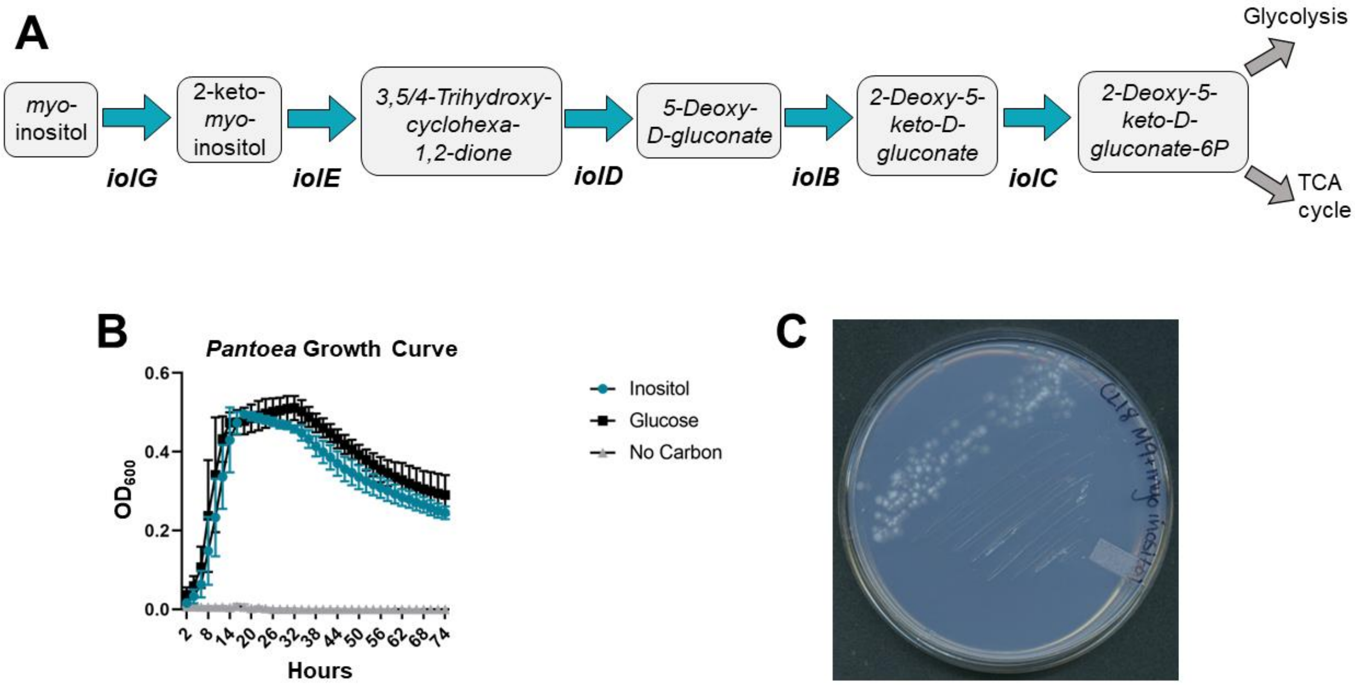


**Figure S3. *Pantoea sp*. R4 and *Streptomyces sp*. CL18 can use inositol as a sole carbon source.** A) A diagram showcasing sequential steps in the bacterial inositol catabolic pathway (*iol* pathway). Rounded rectangles represent intermediary compounds. Enzyme names are listed below arrows. Both *Pantoea sp*. R4 and *Streptomyces sp*. CL18 encode homologs of the 5 *iol* enzymes shown here (iol*GEDBC*). B) Growth dynamics of *Pantoea sp*. R4 in 1x M9 Minimal Salts media with either glucose or inositol as a sole carbon source. A no-carbon control is included. Each curve represents 2 biological replicates with 5 technical replicates each (n=10 wells). Bars represent the standard deviation. C) *Streptomyces sp*. CL18 forms robust colonies on an agar plate with 1x M9 Minimal Salts media with inositol as a sole carbon source. We cannot use optical density to track the growth of *Streptomyces sp*. CL18 in broth, as it grows in aggregates rather than a turbid culture.

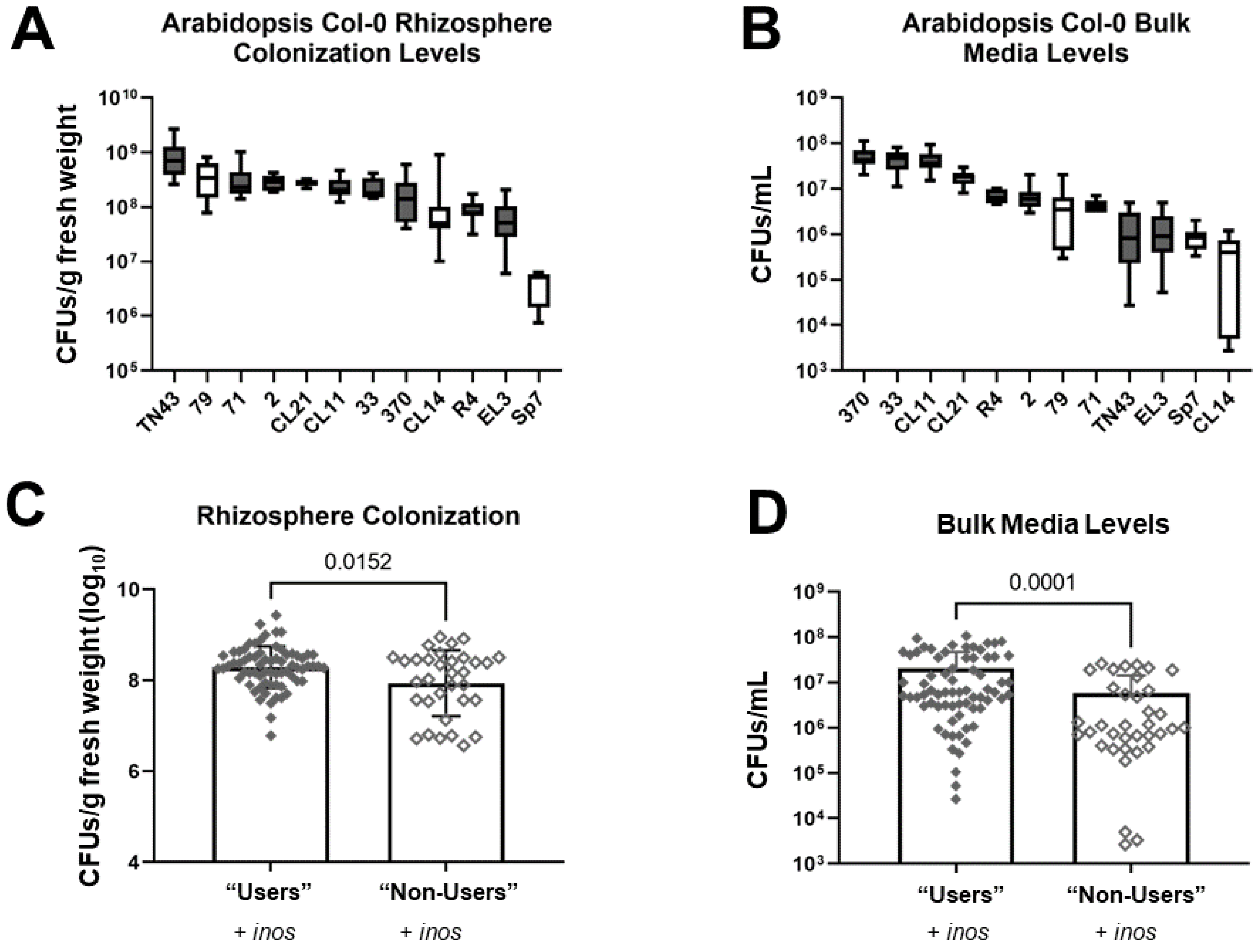


**Figure S4. The pattern linking bacterial inositol catabolism to colonization levels is less robust in the rhizosphere and bulk media.** To further investigate the relationship between inositol catabolism and Arabidopsis colonization, Col-0 seedlings were associated with individual Proteobacterial isolates in low-agar media in magenta jars (MS + inositol). Each jar contained 5 seedlings. After 2 weeks, the 5 seedlings from each jar were pooled and the levels of bacteria in the rhizosphere and bulk media were diluted and plated for CFUs (n=8-11 jars). The rhizosphere colonization levels (A) and bulk media levels (B) are ordered from highest to lowest average CFUs (see Main Figure 3A for paired root colonization levels). For panels A and B, isolates that catabolize inositol have gray box-and-whisker plots, while those that cannot use inositol as a sole carbon source (79, CL21, CL14, and Sp7) have white plots. Whiskers represent min to max values. In contrast to root colonization levels (Main Figure 3), the non-users do not cluster together at the lowest levels for rhizosphere or bulk samples. Panel C shows the individual data points from panel A, aggregated into binary groups according to inositol usage (filled diamonds = "users", open diamonds = "non-users"). Panel D similarly represents the aggregated data from panel B. While inositol "users" have significantly higher average levels in both the rhizosphere and bulk fractions, the magnitude of difference is much smaller than for the roots (roots = 31.1x higher; rhizosphere =1.6x higher; bulk = 3.9x higher). Data points in panel C fit a lognormal distribution and were therefore log-transformed prior to performing an unpaired t-test with Welch's correction. Data points in panel D were analyzed using a non-parametric Mann-Whitney test. Bars in panels C and D represent mean +/- SD. Inositol treatment (+/- inos) is indicated in panels C and D.

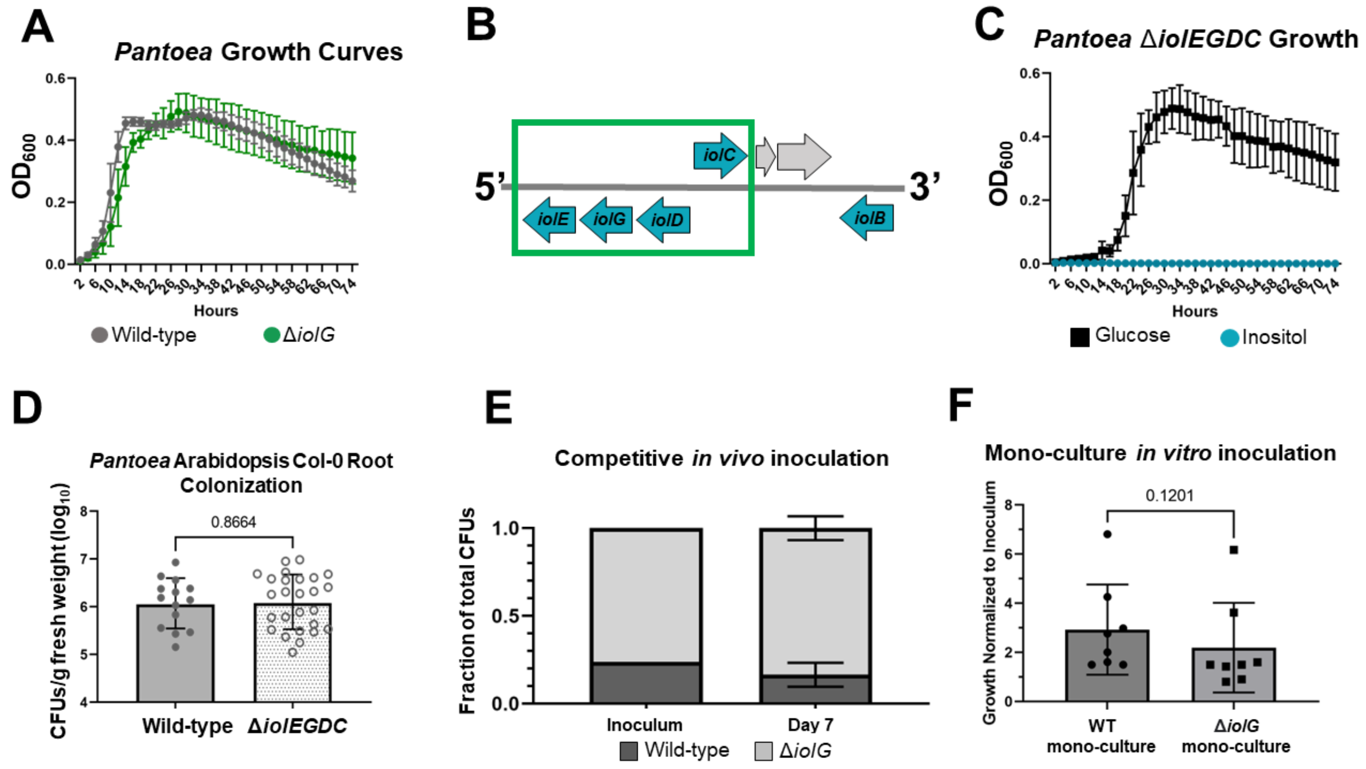


**Figure S5. Single- and multi-gene deletions in the inositol catabolic pathway confer similar *in vitro* and *in vivo* phenotypes.** A) *Pantoea sp*. R4 Δ*iolG* grows similarly to the parental strain on glucose. Each curve represents 2 biological replicates with 6 technical replicates each (n=12 wells). Bars represent the standard deviation. B) A schematic of the genomic region containing the inositol catabolic (*iol)* pathway in *Pantoea sp*. R4's genome. Teal arrows represent genes in the *iol* pathway. Gray arrows represent other genes. A green box surrounds the region deleted for the Δ*iolEGDC* mutant, which contains 4 of the 5 main *iol* genes. JGI gene identifiers for the 4 deleted genes are as follows: *iolG* = 2824863953; *iolE* = 2824863952; *iolD* = 2824863954; and *iolC* = 2824863955. C) *Pantoea sp*. R4 Δ*iolEGDC* can no longer grow on inositol as a sole carbon source, but still grows on glucose. Each curve represents 2 biological replicates with 6 technical replicates each (n=12 wells). Bars represent the standard deviation. D) Similar to the single-gene mutant (Δ*iolG*)*,* the multi-gene mutant *(*Δ*iolEGDC*) colonizes Arabidopsis Col-0 roots at levels similar to the parental strain. Each symbol represents the pooled roots of 3 seedlings. Data points in panel D fit a lognormal distribution and were therefore log-transformed prior to performing an unpaired t-test with Welch's correction. Bars = geometric mean +/- geometric SD. E) *Pantoea sp*. R4 WT and Δ*iolG* strains were inoculated on Col-0 seedlings at the ratio shown in 'Inoculum'. After 7 days, roots were harvested and CFUs from each strain were enumerated. The ratio of each strain in relation to total CFUs was calculated. All replicates received the same inoculum ratio (no error bars), and the ratio after seven days reflected the initial inoculum levels. Data in panel E is from a single biological replicate that included 4 separate growth assays, each with 3 seedlings. A two-way ANOVA (with Šidák's multiple comparisons test and a single pooled variance) revealed no significant differences between Inoculum and Day 7 levels of WT and Δ*iolG* (p-value < 0.05). Bars represent the mean +/- SD. See also Figure 4C for an additional biological replicate with similar outcomes. F) The WT and Δ*iolG* strain were individually inoculated into a root homogenate growth substrate. After 24 hours, cells were counted and normalized to the cell levels in the respective inoculum. The growth (CFUs at 24 hours divided by CFUs in inoculum) of the Δ*iolG* strain was not significantly different than the growth of the parental strain. Each symbol in panel F represents a single well in a 96-well plate. Data points were normalized by dividing the cell numbers after 24 hours by the original inoculum cell numbers. Data in panel F was analyzed with a Mann-Whitney test, and bars = mean +/- SD.
